# Single-cell RNA-sequencing of *Nicotiana attenuata* petal cells reveals the entire biosynthetic pathway of a floral scent

**DOI:** 10.1101/2021.06.28.450226

**Authors:** Moonyoung Kang, Yuri Choi, Hyeonjin Kim, Sang-Gyu Kim

## Abstract

High-throughput single-cell RNA sequencing (scRNA-seq) identifies distinct cell populations based on cell-to-cell heterogeneity in gene expression. By examining the distribution of the density of gene expression profiles, the metabolic features of each cell population can be observed. Here, we employ the scRNA-seq technique to reveal the entire biosynthetic pathway of a flower volatile. The corolla (petals) of the wild tobacco *Nicotiana attenuata* emits a bouquet of scents that are composed mainly of benzylacetone (BA), a rare floral volatile. Protoplasts from the *N. attenuata* corolla were isolated at three different time points, and the transcript levels of >16,000 genes were analyzed in 3,756 single cells. We performed unsupervised clustering analysis to determine which cell clusters were involved in BA biosynthesis. The biosynthetic pathway of BA was uncovered by analyzing gene co-expression in scRNA-seq datasets and by silencing candidate genes in the corolla. In conclusion, the high-resolution spatiotemporal atlas of gene expression provided by scRNA-seq reveals the molecular features underlying cell-type-specific metabolism in a plant.

## Introduction

Flowers produce diverse volatiles for various ecological interactions. To date, more than 1,700 different floral scents have been identified (Knudsen et al., 2006): these scents are classified as terpenoids, fatty acid-derivatives, and phenylpropanoid/benzenoids, according to their biosynthetic pathways (Muhlemann et al., 2014; Knudsen and Gershenzon, 2020). The major molecular mechanisms underlying floral scent diversity are the duplication and subsequent mutation (neofunctionalization) of biosynthetic genes (Moghe and Last, 2015). A mutation in a coding region can generate new volatile compounds by binding to the original substrate or to similar substrates. For example, a single amino-acid substitution in *Oryza sativa* terpene synthase 1 (TPS1) changes the ratio between two products: (*E*)-β-caryophyllene and germacrene A (Chen et al., 2014). Mutations in a cis-regulatory element may trigger changes in the spatiotemporal expression of biosynthetic enzymes, resulting in the generation of a new metabolic pathway. Because small changes in enzymes can result in new compounds, identifying the genes responsible for floral scent biosynthesis requires a multidisciplinary approach that includes functional prediction based on sequence homology to characterized enzymes, biochemical assay, and *in vivo* gene manipulation.

To identify the genes responsible for floral scent production, DNA/RNA sequencing techniques, microarrays, and several statistical methods have been widely used. Candidate genes have been prioritized by investigating correlations among transcript abundances and target compounds (Magnard et al., 2015; Xu et al., 2018; Verdonk et al., 2005). By analyzing differential gene expression among the rose cultivars that vary widely in floral scent emission, researchers identified a sesquiterpene synthase (Guterman et al., 2002) and a nudix hydrolase (Magnard et al., 2015), a new type of monoterpene synthase. In addition, when a biosynthetic gene in a multi-step pathway is available, this gene can be used as “bait” to calculate its degree of association to the entire transcriptome (gene co-expression network analysis) (Wisecaver et al., 2017) and to construct a gene cluster that would regulate the production of metabolites, including volatiles (Li et al., 2020). For instance, TPS11 enzymes in *Arabidopsis thaliana* flowers synthesize (+)-α-barbatene and (+)-thujopsene. These volatiles are further oxidized by CYP706A3, which was discovered through co-expression analysis with TPS11 (Boachon et al., 2019). These results show that the successful identification of biosynthetic genes of specialized volatiles through gene-to-metabolite or gene-to-gene correlation requires high diversity in expression that comes from underlying genetic, environmental, time-series or tissue-specific diversity in the material analyzed (Wong et al., 2020).

Flower scents are mainly synthesized in the petals. In particular, several volatile biosynthetic genes are detected in the epidermal cells of petals, for example: (*S*)-linalool synthase (Dudareva et al., 1996) and isoeugenol *O*-methyltransferase (Dudareva and Pichersky, 2000) in *Clarkia breweri* flowers; benzoic acid carboxyl methyltransferase in *Antirrhinum majus* flowers (Kolosova et al., 2001); and orcinol *O*-methyltransferase in rose cultivars (Scalliet et al., 2006). Separating these producing cells from the rest of the cells in the entire petal can increase the accuracy of a co-expression analysis; laser microdissection and marker-based cell sorting methods have been used to isolate targeted cells (Wong et al., 2020). Recently, several high-throughput single-cell RNA sequencing (scRNA-seq) techniques have emerged for analyzing transcriptomes expressed in single cells isolated from tissues or organs. The scRNA-seq database shows cellular heterogeneity and molecular processes specific to cell types in plant tissues, such as roots (Ryu et al., 2019; Denyer et al., 2019; Jean-Baptiste et al., 2019; Shulse et al., 2019), leaves (Kim et al., 2021; Lopez-Anido et al., 2020), and shoot apical meristems (Satterlee et al., 2021; Zhang et al., 2021). Assuming that all biosynthetic genes of a given floral scent are expressed in a single cell and that the regulation of those genes is coordinated, the scRNA-seq database of petals can be useful in prioritizing candidate genes, as it allows researchers to calculate correlations among genes expressed in several thousands of petal cells (Fridman and Pichersky, 2005; Wong et al., 2020).

Flowers in the wild tobacco *Nicotiana attenuata* emit benzylacetone (BA, 4-phenyl-2-butanone) at night to attract pollinators, such as *Manduca sexta* hawkmoths (Baldwin, 1997; Haverkamp et al., 2016; Euler and Baldwin, 1996), and to prevent floral damage from cucumber beetles (Kessler et al., 2019). The minor floral scent compounds are benzyl alcohol, cis-3-hexenol, and (*E*)-α-bergamotene (Kessler and Baldwin, 2007). Most BA is produced from corolla limbs (Kessler and Baldwin, 2007; Euler and Baldwin, 1996), which is regulated by the circadian clock (Yon et al., 2016). L-phenylalanine (Phe) is a precursor of BA, and phenylalanine ammonia-lyase 4 (PAL4) is known to be the first enzyme in the BA biosynthetic pathway (Guo et al., 2020). In the plant kingdom, the major Phe-derived floral volatiles are phenylpropanoid (a phenyl ring with a three-carbon side chain, C6-C3) and benzenoid (C6-C1) (Widhalm and Dudareva, 2015). However, BA possesses a rare structure, namely, one with a C6-C4 structure (phenylbutanoids). Here, we performed the scRNA-seq of protoplasts isolated from *N. attenuata* corolla and defined distinct cell clusters based on the cell-to-cell variation in gene expression. Gene co-expression analysis with scRNA-seq datasets enables us to identify the entire set of BA biosynthetic genes.

## Results

### Single-cell RNA sequencing defines distinct cell types in the corolla

To generate a single-cell transcriptome atlas of genes involved in BA biosynthesis, we isolated protoplasts from *N. attenuata* corolla limbs and throats at three time points (Zeitgeber Time 8, ZT 12, and ZT 16) when BA emission increases (Fig. 1A). The protocol of protoplast isolation was optimized to retain cell integrity and the temporal transcriptome; the whole process, from flower excision to generating single-cell gel bead-in-emulsion, took about 2 h. The 10X Genomics Chromium and the Illumina sequencing platforms were used to generate scRNA-seq libraries. A total of 16,465 genes from 3,756 cells and approximately 1,300 genes per cell were detected. To evaluate the processes involved in scRNA-seq, we generated a pseudo-bulk dataset for each time point by pooling scRNA-seq unique molecular identifiers (UMIs) and performed conventional bulk RNA sequencing (bulk RNA-seq) with corolla limbs collected at the same time points (ZT 8, ZT 12, and ZT 16); these pseudo-bulk datasets were compared with the bulk RNA-seq datasets. The global transcriptome profiles of two datasets were highly correlated (Pearson’s correlation, r >0.8, Fig. S1). We removed stress-responsive genes that were highly induced in scRNA-seq libraries compared to bulk RNA-seq datasets (Table S1).

**Figure 1.**
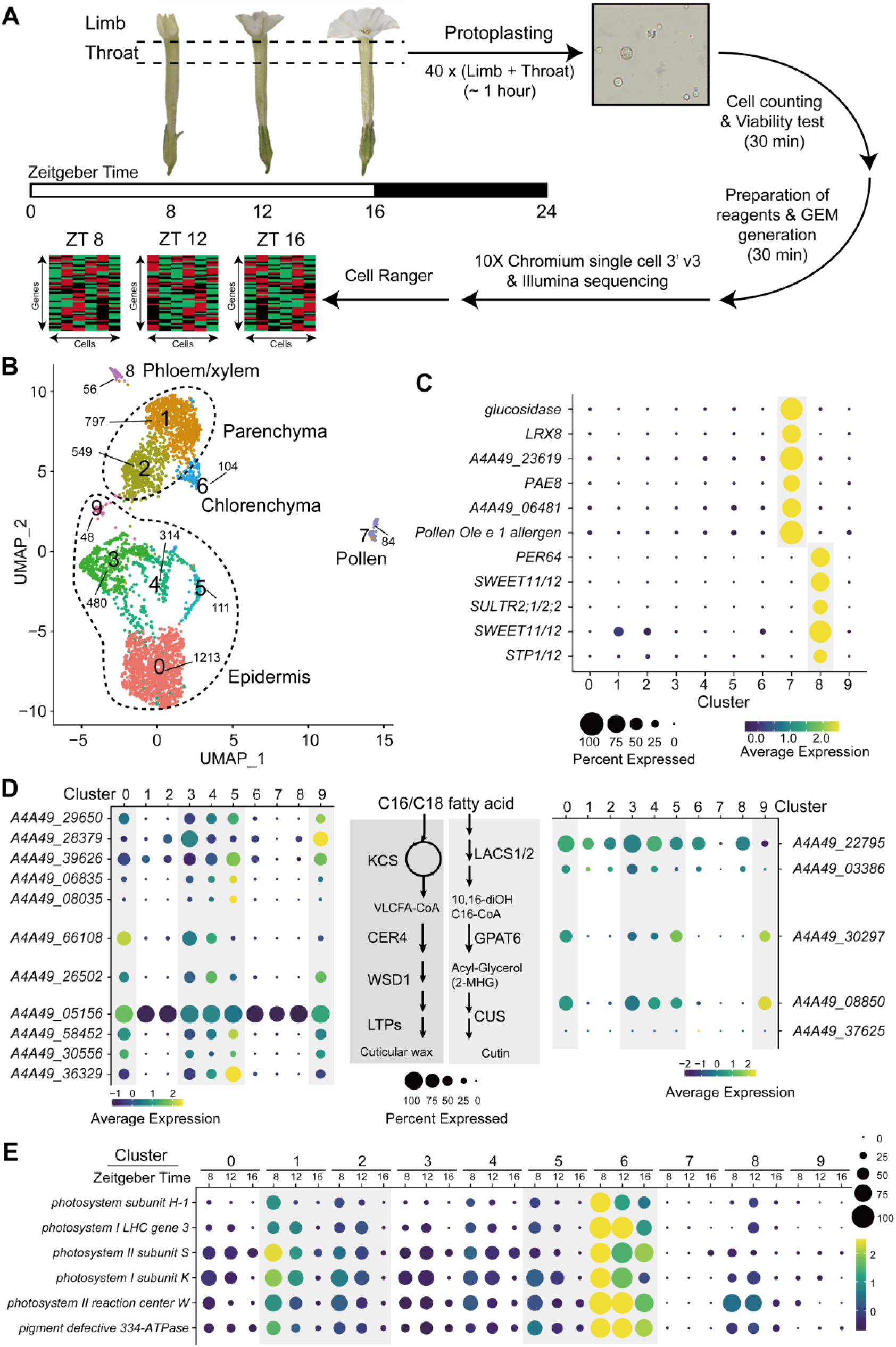
Single-Cell RNA sequencing characterizes distinct *N. attenuata* corolla cell types. (A) Corolla limbs and throat cups were sampled from 40 *N. attenuata* flowers at each Zeitgeber Time (ZT), and scRNA-seq libraries were generated. The total experimental procedures were finished within 2 h to reduce the effect of protoplasting. (B) A uniform manifold approximation and projection (UMAP) plot of the 3,756 cells classified into 5 major cell types, composed of total 10 distinct clusters. The number of cells of each cluster is given next to clusters. (C) Pollen marker genes and phloem/xylem marker genes are exclusively detected in clusters 7 and 8, respectively. SULTR2;1/2;2, a sulfate transporter, and SWEET11/12 and a sugar transporter1/12 (STP1/12) are known to be exclusively expressed in *A. thaliana* vasculatures. All marker genes in cluster 7 are specifically expressed in pollen or pollen tubes (Brockmöller et al., 2017). Dot size and color indicate the percentage of cells expressing a marker gene and mean expression value across cells in each cluster, respectively. (D) Most biosynthetic genes of cuticular wax and cutin are highly enriched in clusters 0, 3, 4, 5, and 9. These genes are known to be expressed in the epidermis. CER4, ECERIFERUM4; CUS, cutin synthase-like; GPAT6, glycerol-3-phosphate acyltransferase 6; KCS, 3-ketoacyl-CoA synthase; LACS1/2, long-chain acyl-CoA synthase 1/1; LTP, lipid-transfer proteins; VLCFA, very-long-chain fatty acids; WSD1, wax ester synthase/diacylglycerol acyltransferase 1. (E) Photosynthesis-associated genes are highly expressed in cluster 6 at all time points and in clusters 1, 2, and 6 at ZT 8. These marker genes are known to be expressed in the internal layers of plant tissue.

We next categorized single cells into clusters based on their shared transcriptomic profiles using the Seurat package (Satija et al., 2015; Butler et al., 2018). Cells were projected and visualized on a uniform manifold approximation and projection (UMAP) plot, which showed 10 distinct clusters in two-dimensional space (Fig. 1B). Because there were no well-known marker genes for petal cells, we assigned cell identities based on the putative biochemical functions of cluster-specific genes. The *N. attenuata* transcriptome was annotated based on the homology of protein-coding sequences to previously investigated genes in several model plants (Brockmöller et al., 2017). In cluster 7, several pectin esterase genes, extensin families, and homologs of pollen allergens were specifically detected; these cells were defined as pollen that had attached to the corolla’s surface (Fig. 1C). In cluster 8, we detected several orthologs of *AtSWEET11*/*12* and *AtSULTR2;1* genes, which are known to be specifically expressed in phloem and xylem (Fig. 1C) (Chen et al., 2012; Kim et al., 2021; Maruyama-Nakashita et al., 2015).

We next assigned the remaining major clusters to three groups of cells: epidermis, chlorenchyma, and parenchyma. Epidermal cells are known to synthesize cuticular wax and cutin for their cuticle layer. Several genes involved in cuticular wax biosynthesis were specifically enriched in clusters 0, 3, 4, 5, and 9; *ketoacyl-CoA synthase, ECERIFERUM4, wax ester synthase/diacylglycerol acyltransferase 1*, and several *lipid-transfer proteins*, which may participate in wax synthesis or transport, were highly expressed (Figs. 1D, S2). We also found that clusters 0, 3, 4, 5, and 9 contained higher levels of cutin biosynthetic genes (*glycerol-3-phosphate acyltransferase 6* and *cutin synthase-like*) compared to other clusters. These results suggest that clusters 0, 3, 4, 5, and 9 are epidermal cells. Cluster 6 was putatively defined as a chlorenchyma cell, because complex subunits of photosystem I and II were highly expressed in this cluster (Figs. 1E, S3). As shown in Fig. 1A, the throat cup of the corolla retained chloroplasts. Photosynthetic genes are well-known to have a clear diurnal rhythm, with high levels during the day and low levels during the night (Kim et al., 2011). Our scRNA-seq data showed that the transcript levels of several photosynthetic genes gradually decreased from ZT 8 (in the middle of the day) to ZT 16 (the transition time from light to dark) (Fig. 1E). Clusters 1 and 2 had low levels of cuticle biosynthetic genes and relatively high levels of photosynthetic genes. These results suggest that in the corolla limb, clusters 1 and 2 are parenchyma cells.

### Correlation analysis at single-cell resolution prioritizes candidate genes for BA biosynthesis

In the corolla, the initial step of BA biosynthesis is the non-oxidative deamination of Phe by the action of *Nicotiana attenuata* PAL4 (NaPAL4, Fig. 2A) (Guo et al., 2020). The product of NaPAL4 enzyme, *t*-cinnamic acid, is likely converted to cinnamoyl-Coenzyme A (CoA) by a homologous protein of 4-coumarate:CoA ligase (4CL) or cinnamate:CoA ligase (CNL) (Klempien et al., 2012) without further hydroxylation, because BA has no hydroxyl group on the aromatic ring. Next, the cinnamoyl-CoA is likely condensed with one molecule of malonyl-CoA, which yields benzalacetone (4-phenyl-3-buten-2-one), a C6-C4 moiety of phenylbutanoid. Abe *et al*. found that a novel polyketide synthase named benzalacetone synthase produces 4-hydroxybenzalacetone (4-(4-hydroxyphenyl)-3-buten-2-one) from 4-coumaroyl-CoA and one molecule of malonyl-CoA (Abe et al., 2001; Morita et al., 2010; Shimokawa et al., 2012). *N. attenuata* contains four polyketide synthase enzymes (NaPKS1/2/3/4, renamed from NaCHAL1/2/3/4, respectively) homologous to chalcone synthase or benzalacetone synthase (Fig. S4) (Abe et al., 2001); silencing *NaPKS3* (A4A49_39367) dramatically reduces the level of BA emission (Guo et al., 2020; Kessler et al., 2008). However, in the target region of the previous RNAi construct, *NaPKS3* sequence differed from *NaPKS1* (A4A49_08280), *NaPKS2* (A4A49_34074), and *NaPKS4* (A4A49_65921) by only 34, 10, and 39 nucleotides, respectively (Kessler et al., 2008) (Fig. S5). Therefore, we hypothesized that one or some of NaPKSs play a role to convert cinnamoyl-CoA to benzalacetone. Finally, benzalacetone appears to be reduced by an unknown reductase, which leads to the production of BA.

**Figure 2.**
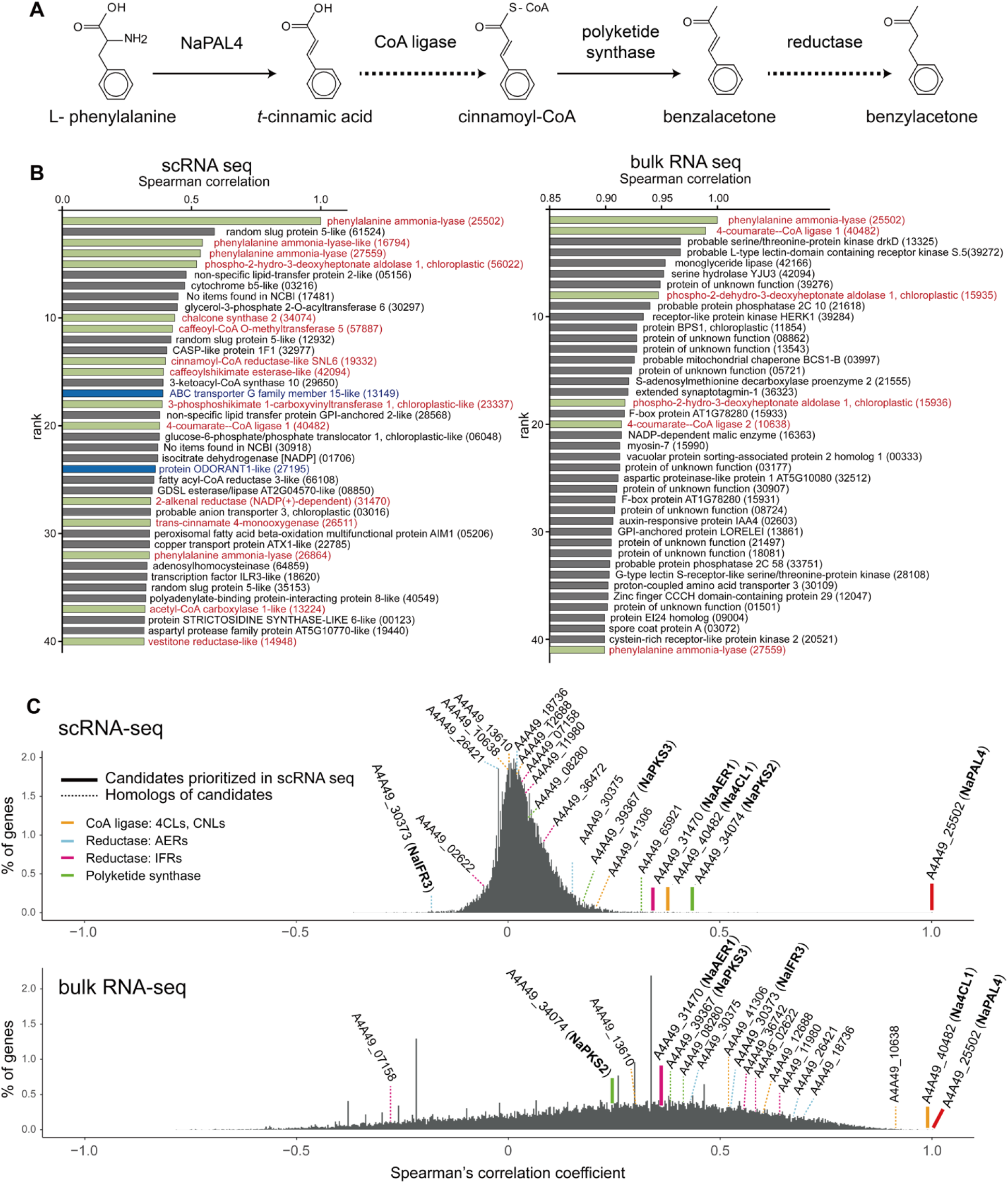
Identification of benzylacetone biosynthetic genes via correlation analysis at single-cell resolution. (A) Benzylacetone (BA) biosynthetic pathway proposed based on the biochemical function of NaPAL4 and NaPKS1/2/3/4 (solid lines), which have been characterized in several literatures. Dashed lines indicate a putative reaction mediated by unknown enzymes: CoA ligase, and reductase. NaPAL4, *Nicotiana attenuata* phenylalanine ammonia-lyase 4; NaPKS, *N. attenuata* polyketide synthase. (B) Two correlation analyses display different top-ranked gene profiles. Genes are ranked by their correlation coefficients with NaPAL4. ScRNA-seq based analysis (left panel) generated more relevant profile where several phenylpropanoid pathway associated genes are included; green bar: putative phenylpropanoid pathway genes, blue bar: homologs of transporters and transcription factor studied in petunia. (C) Percentage of genes (y-axis) that have a given Spearman’s rank correlation coefficient (x-axis) with *NaPAL4* expression in scRNA-seq and bulk RNA-seq. Genes encoding CoA ligase, reductases, and chalcone synthases appear in different colors on the histograms. Thick lines highlight the CoA ligase, reductase, and chalcone synthase most correlated with *NaPAL4* in scRNA-seq. NaAER1, *N. attenuata* 2-alkenal reductases 1; NaCNL, *N. attenuata* cinnamate:CoA ligase; NaIFR, *N. attenuata* isoflavone reductase; Na4CL1, *N. attenuata* 4-coumarate:CoA ligase.

To identify BA biosynthetic genes, the CoA ligase, the PKS, and the reductase, we hypothesized that all BA biosynthetic genes would be co-regulated over time at the single-cell level. We calculated the Spearman’s rank correlation coefficient of *NaPAL4* expression with the whole gene expression detected in our scRNA-seq data and in previously published bulk RNA-seq data from tissue taken from more than twenty plant parts, including seed, leaf, root, stem, anther, style, stigma, ovary, nectary, pedicel, and corolla (Brockmöller et al., 2017; Li et al., 2016). We then compared functional relationships among the top 40 ranked genes with *NaPAL4* in both datasets (Fig. 2B): genes involved in phenylpropanoid metabolism were detected more in scRNA-seq (13 among 40 genes) than in bulk RNA-seq (6 among 40 genes). The top 40 ranked genes in scRNA-seq and bulk RNA-seq data contained one putative 4CL/CNL-like gene (Na4CL1, A4A49_40482), suggesting that Na4CL1 was the strongest candidate enzyme for producing BA.

We also found that prioritized candidates and their homologs were distributed differently in scRNA-seq and bulk RNA-seq datasets. Among four *NaPKSs, NaPKS2* was the gene that ranked highest in scRNA-seq datasets, followed by *NaPKS3* (Fig. 2C). However, in bulk RNA-seq datasets *NaPKS3* was correlated more with *NaPAL4* than did *NaPKS2* (Fig. 2C). Five homologues of AER and four homologues of isoflavone reductase (IFR), another family of reductase, were found in the scRNA-seq and bulk RNA-seq datasets. Interestingly, one putative 2-alkenal reductase (NaAER1, A4A49_31470) was highly ranked only in scRNA-seq, and no reductase was found among the top 1,800 ranked genes in the bulk RNA-seq (Figs. 2B, C). In the bulk RNA-seq, all *IFR* genes and three of five *AER* homologues were more highly correlated with *NaPAL4* expression than did the *NaAER1* (Fig. 2C). The distribution of the correlation coefficient in scRNA-seq differed from its distribution in bulk RNA-seq. A large number of genes in scRNA-seq appeared to have a low correlation with *NaPAL4*: 99.46 % of genes had a correlation coefficient between -0.25 and 0.25. The histogram from bulk RNA-seq showed a broad and skewed distribution of correlation coefficients.

### Na4CL1, NaPKS2, and NaAER1 are involved in BA biosynthesis

To test the model for BA biosynthesis that emerged from the co-expression analysis in scRNA-seq datasets, we firstly silenced the expression of *Na4CL1* by virus-induced gene silencing (VIGS) and measured the level of floral volatiles in the headspace and in endogenous cells. Silencing *Na4CL1* reduced levels of BA both in flower headspace (Fig. 3A) and in endogenous cells (Fig. 3B). It is noteworthy that other *Na4CL1* homologues – A4A49_41306, A4A49_10638 – and *NaCNL* homologues – A4A49_13610, and A4A49_12688 – which are expressed in *N. attenuata* flowers were not silenced in *Na4CL1*-silenced corolla limb (Figs. 3C, S6). To evaluate the catalytic ability of Na4CL1, purified Na4CL1 proteins were tested with *t*-cinnamic acid and several C6-C3 compounds with hydroxyl and methoxy group (4-coumaric acid, caffeic acid, and ferulic acid) in the same conditions (Fig. 3D). Best-fit kinetic values were calculated by fitting with Michaelis Menten curve. Na4CL1 showed ability to catalyze the formation of CoA-thioesters of *t*-cinnamic acid and similar C6-C3 compounds. It suggests that Na4CL1 can activate carboxylate groups in several C6-C3 compounds in *N. attenuata*, mediating reactions on carbon chain of phenylpropanoids. Determined *K*_*m*_ values showed that Na4CL1 has higher affinity to *t*-cinnamic acid and 4-coumaric acid than caffeic acid and ferulic acid.

**Figure 3.**
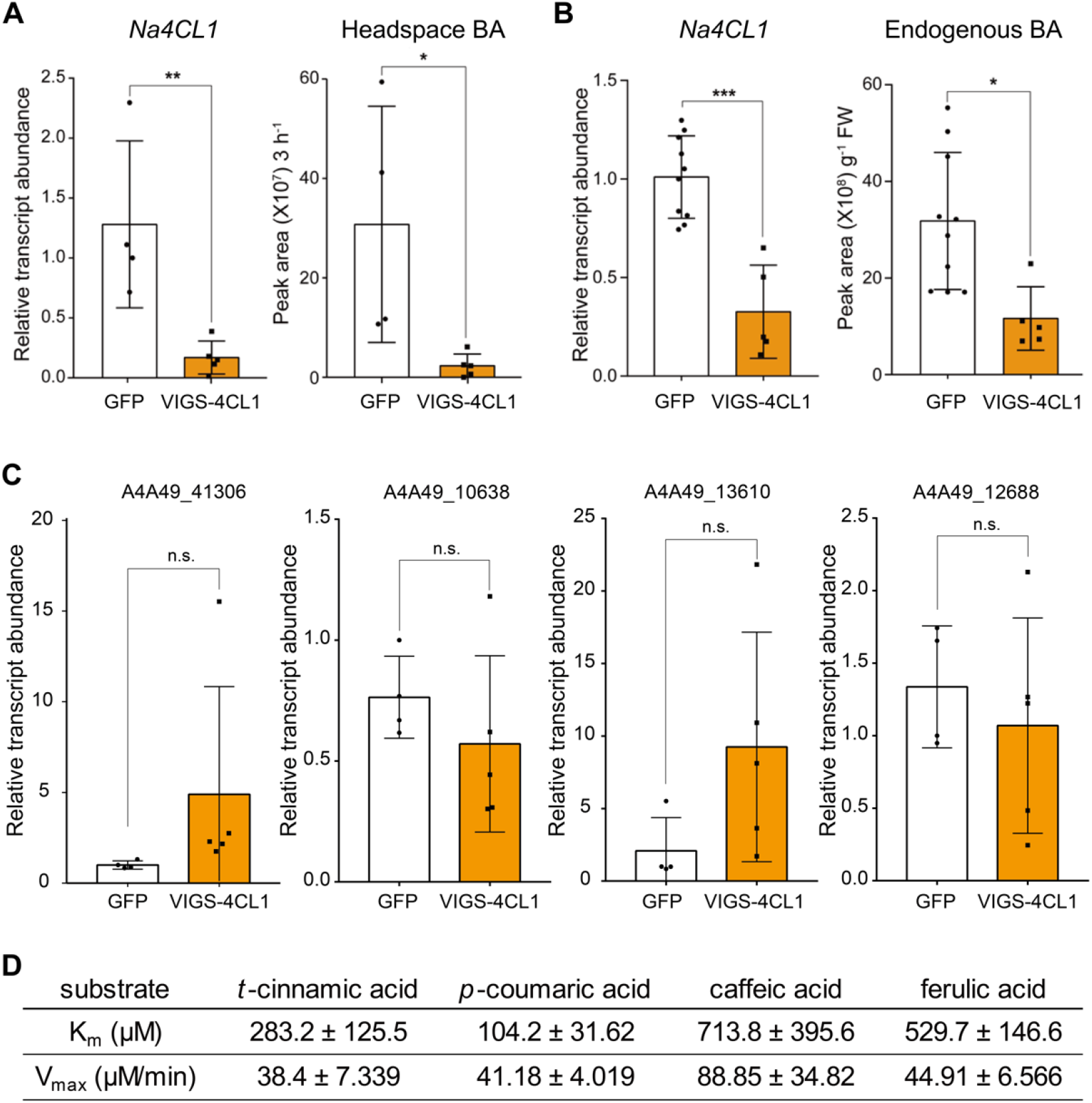
Na4CL1 participates in benzylacetone biosynthesis in *N. attenuata* flowers. (A, B) Mean (± SD) levels of headspace benzylacetone (BA) and endogenous BA were measured in *Na4CL1*-silenced flowers (mean ± SD; *, *p*<0.05; **, *p*<0.01; ***, *p*<0.001; two-tailed Student’s t-test). GFP, plant infected TRV2:GFP (a truncated form), which was used as a negative control; VIGS-4CL1, a *Na4CL1*-silenced plant, which was generated by virus-induced gene silencing (VIGS). (C) Transcript accumulation of four *Na4CL1* homologues in *Na4CL1*-silenced flowers. (D) Kinetic parameters of Na4CL1 determined from *in vitro* enzyme assay of heterologously purified Na4CL1 protein. Best-fit catalytic values were calculated by fitting with Michaelis Menten curve, showing that Na4CL1 can catalyze the formation of CoA-thioesters of *t*-cinnamic acid and similar C6-C3 compounds. Na4CL1, *Nicotiana attenuata* 4-coumarate:CoA ligase.

We next tried to verify whether NaPKS3 is involved in BA biosynthesis as previously suspected (Guo et al., 2020), because the analysis of scRNA-seq datasets showed that *NaPKS2* was more correlated with *NaPAL4* than did *NaPKS3*. Although we designed the VIGS construct as specific as possible, we observed the co-silencing of *NaPKS1/2/3/4* (Figs. 4A, S7) in *NaPKS3*-silenced plants; high sequence similarity was observed among *NaPKSs* (Fig. S5). As previously reported, silencing *NaPKSs* reduced levels of BA in headspace (Fig. 4B) and corolla limb (Figs. 4C, D). Although both *NaPKS2* and *NaPKS3* were strongly expressed in flower tissues (Fig. S8), scRNA-seq datasets indicated that *NaPAL4* was more correlated with *NaPKS2* than *NaPKS3* (Fig. 2C). To clarify which NaPKSs are required for BA biosynthesis, we incubated the same amount of purified NaPKS2 or NaPKS3 proteins with Na4CL1 products (cinnamoyl-CoA) and malonyl-CoA (as carbon donor). The result showed that the only NaPKS2-containing mixture produced benzalacetone (Fig. 4E). We did not detect benzalacetone in the NaPKS3-containing mixture (Fig. 4E). The result suggests that NaPKS2, rather than NaPKS3, plays a major role for BA biosynthesis, as predicted by scRNA-seq correlation analysis (Fig. 2C).

**Figure 4.**
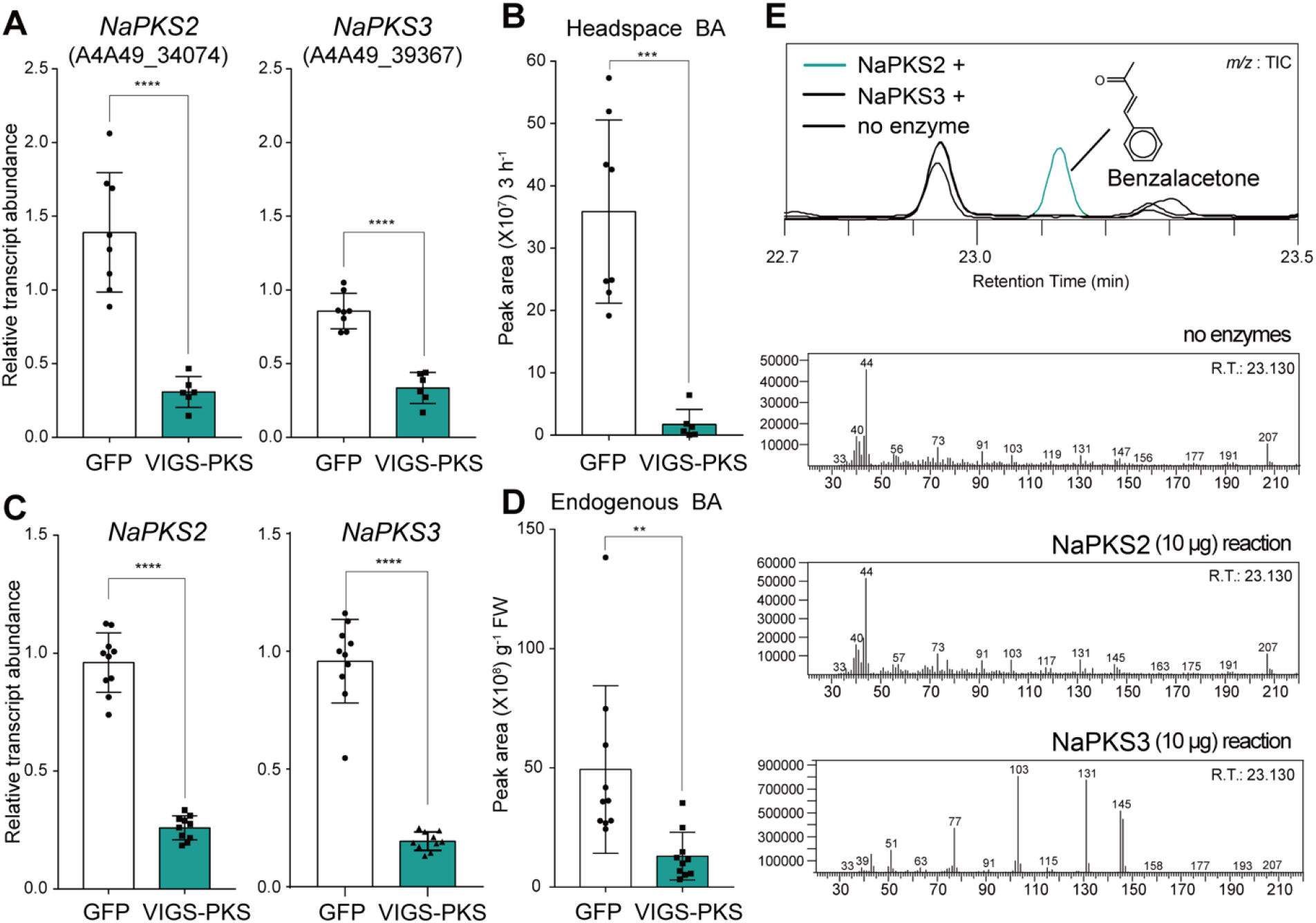
NaPKS2 is the enzyme that participates in benzylacetone biosynthesis in *N. attenuate* flowers. (A) *NaPKS2* and *NaPKS3* are co-silenced in *NaPKS3*-silenced plants. GFP, plant infected TRV2:GFP (a truncated form), which was used as a negative control; VIGS-PKS, a *NaPKSs* co-silenced plant, which was generated by virus-induced gene silencing (VIGS). (mean ± SD; **, *p*<0.01; ***, *p*<0.001; ****, *p*<0.0001; two-tailed Student’s t-test). NaPKS, *Nicotiana attenuata* polyketide synthase. (B) Mean (± SD) levels of headspace benzylacetone (BA) show that *NaPKS2*/*3*-silenced flowers emit less BA. (C, D) *NaPKS2*/*3*-silenced flowers harbor less endogenous BA in tissue. (E) *In vitro* enzyme assay reveals that NaPKS2, not NaPKS3, can synthesize benzalacetone, a putative intermediate of BA biosynthesis. A novel peak detected in NaPKS2-contained mixtures represents benzalacetone, based on its fragmentation pattern.

To identify the last step enzyme of BA biosynthesis, we silenced the candidate reductase *NaAER1* by VIGS and measured the level of floral volatiles in the headspace and in endogenous cells. Silencing *NaAER1* also impaired BA biosynthesis in flowers without silencing other homologues; the levels of BA emission and endogenous BA were reduced in *NaAER1*-silenced flowers (Figs. 5A, B, S9). We detected a novel compound emitted from *NaAER1*-silenced flowers (Fig. 5C). Metabolite fragmentation patterns and standard compound injections verified that this compound is benzalacetone. In addition, *in vitro* enzyme assay showed that purified NaAER1 proteins with NADPH, a biological reducing agent, can convert benzalacetone to BA (Fig. 5D). The accumulation of benzalacetone in *NaAER1*-silenced flowers and the *in vitro* assay confirmed that NaAER1 is the final enzyme involved in BA biosynthesis (Fig. 5E).

**Figure 5.**
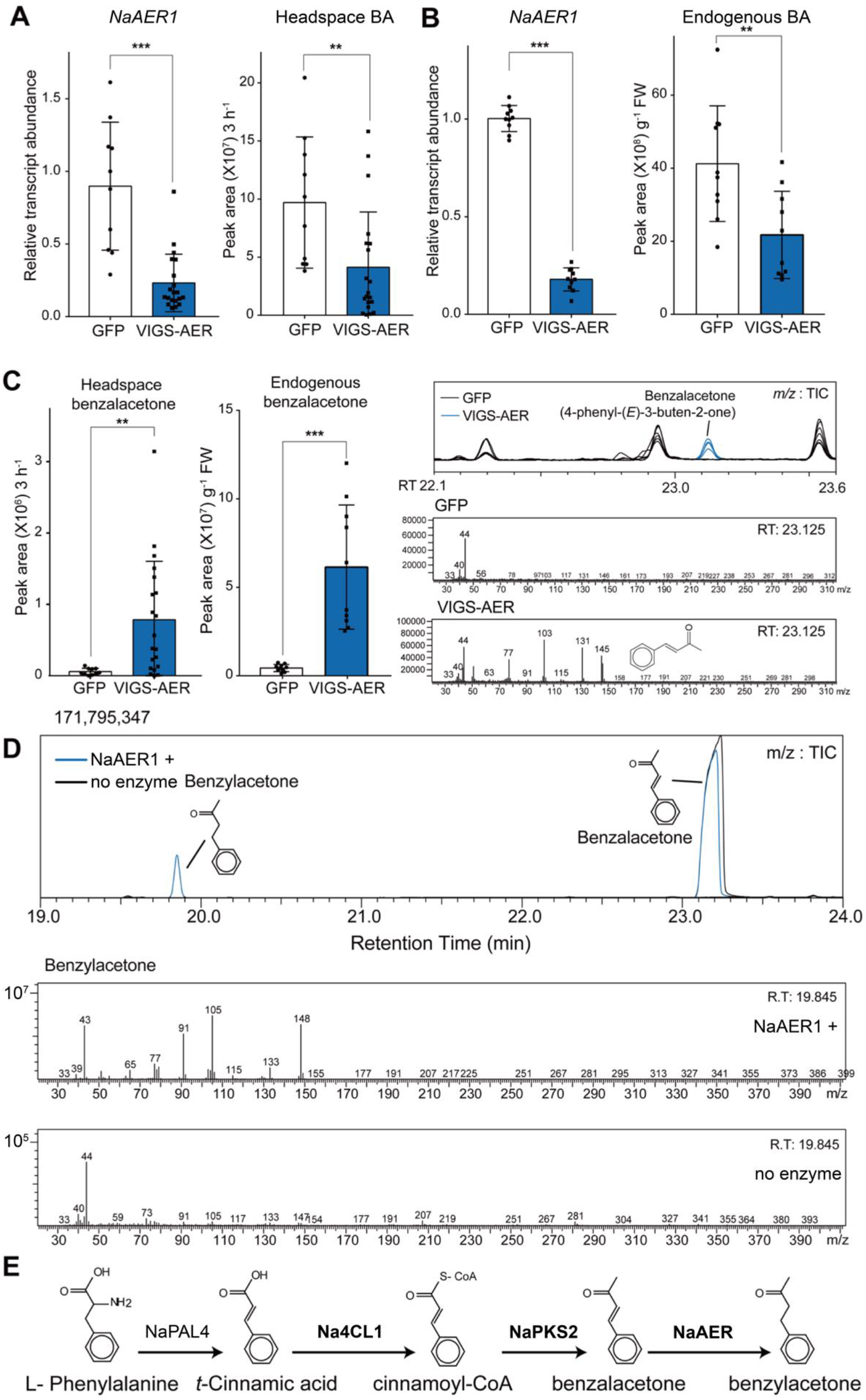
NaAER1 catalyzes the final step in benzylacetone biosynthesis. (A, B) Mean (± SD) levels of headspace benzylacetone (BA) and endogenous BA were measured in *NaAER1*-silenced flowers (mean ± SD; **, *p*<0.01; ***, *p*<0.001; two-tailed Student’s t-test). BA levels were reduced in *NaAER1*-silenced flowers. GFP, plant infected TRV2:GFP (a truncated form), which was used as a negative control; VIGS-AER, a *NaAER1*-silenced plant, which was generated by virus-induced gene silencing (VIGS). (C) Benzalacetone, a metabolic intermediate of BA, was emitted from *NaAER1*-silenced flowers and not from control flowers (mean ± SD; **, *p*<0.01; ***, *p*<0.001; two-tailed Student’s t-test). Right panel shows a novel GC-MS peak in an *NaAER1*-silenced flower, which has the same retention time and mass fragmentation pattern as the benzalacetone standard. (D) *In vitro* assay of NaAER1. NaAER1 reduces the double bond of benzalacetone, completing BA biosynthesis. (E) Complete elucidation of BA biosynthesis pathway.

Recently, Guo *et al*. reported that *NaIFR3* showed a strong correlation with *NaPAL4* in the same bulk RNA-seq dataset used in this study (Guo et al., 2020). Although we also observed the positive correlation between *NaIFR3* and *NaPAL4* in bulk RNA-seq data, there was no positive correlation between these two genes in the scRNA-seq dataset (Fig. 2C). Furthermore, we found that levels of BA emission in *NaIFR3*-silenced flowers did not differ from levels in control flowers (Fig. S10A, no significant difference). To further test these results, we measured the endogenous levels of BA in *NaIFR3*-silenced flowers. No difference in endogenous BA was found between control and *NaIFR3*-silenced flowers (Fig. S10B).

### BA is synthesized in the epidermal layer

Among the 3,756 cells, 1,648 contained at least one molecule of *NaPAL4* mRNA (Fig. S11). The 80 % of *NaPAL4*-expressing cells belonged to epidermal cells (clusters 0, 3, and 4), which suggests that BA is synthesized in the epidermal layer of the corolla. We calculated the number of cells harboring individual gene or combination of *NaPAL4, Na4CL1, NaPKS2*, and *NaAER1* genes. Of 3,756 cells, 1,097 contained all four genes; 94% of these cells were found in clusters 0, 3, and 4 (epidermal cells) (Fig. 6A). Most *NaPAL4*-expressing cells (67%) contained all three genes: *Na4CL1, NaPKS2*, and *NaAER1*. To visualize the spatial distribution of cells in which two target genes were co-expressed, we merged two feature plots of *NaPAL4* and one of *Na4CL1, NaPKS2*, and *NaAER1* expression (Fig. 6B). The results showed that a high level of co-expression (yellowish cells) was mainly found in cluster 0, implicating the scent biosynthesis step occur in the epidermis. Phe biosynthetic genes were detected mainly in clusters 0, 3, and 4, but these genes were also expressed in other clusters (Fig. 6C). Dotplots showed that *NaPAL2*/*3*/*4* with *Na4CL1, NaPKS2*, and *NaAER1* were highly expressed in the epidermal cells (Fig. 6C).

**Figure 6.**
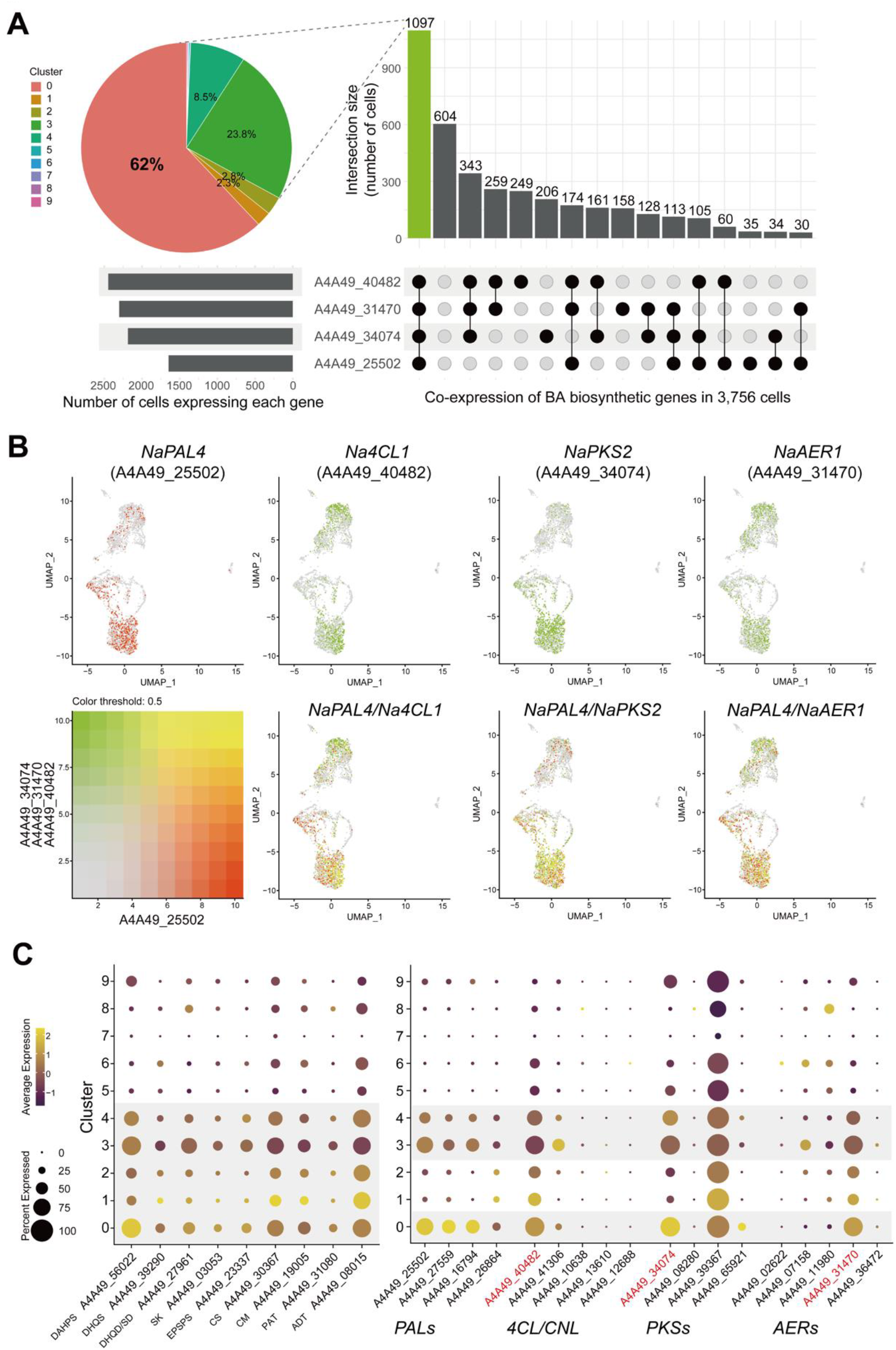
Characterization of benzylacetone biosynthetic cell clusters. (A) An UpSet plot shows numbers of cells harboring individual gene or combination of *NaPAL4, Na4CL1, NaPKS2*, and *NaAER1* genes. A pie chart represents the distribution of clusters of cells expressing all four genes. Clusters 0, 3, and 4 (epidermis) contain 94.3 % of cells co-expressing all four genes. (B) Featureplots visualize co-expression between *NaPAL4* and one of *Na4CL1, NaPKS2*, and *NaAER1*. Cells that coexpress genes mostly locates in cluster 0. (C) Dotplots of Phe biosynthetic genes (primary metabolism) and BA biosynthetic genes (secondary metabolism). Phe is mainly synthesized in clusters 0, 1, 3, and 4, and BA synthesis occurs in clusters 0, 3, and 4. DAHPS, phospho-2-dehydro-3-deoxyheptonate aldolase; DHQS, 3-dehydroquinate synthase; DHQD/SD, bifunctional 3-dehydroquinate dehydratase/shikimate dehydrogenase; SK, shikimate kinase; EPSPS, 3-phosphoshikimate 1-carboxyvinyltransferase 1; CS, chorismate synthase; CM, chorismate mutase; PAT, bifunctional aspartate aminotransferase and glutamate/aspartate-prephenate aminotransferase; ADT, arogenate dehydratase/prephenate dehydratase

### Diurnal rhythms of BA biosynthetic genes and BA emissions

Because *N. attenuata* flowers emit BA rhythmically (Yon et al., 2016), we asked whether all BA biosynthetic genes share this diurnal expression pattern. In our experiments, BA emission began near dusk and peaked at ZT 16 (Fig. 7A). Quantitative PCR analysis showed that the transcript levels of *NaPAL4, Na4CL1, NaPKS2*, and *NaAER1* peaked approximately 4 h before BA emission peaked, suggesting that BA emission is regulated mainly at the transcriptional level. In petunia flowers, rhythmic scent emission is also regulated transcriptionally (Amrad et al., 2016; Dexter et al., 2007; Fenske et al., 2015; Klempien et al., 2012). Our scRNA-seq data also showed similar expression patterns of BA biosynthetic genes in each cluster. For instance, the violin plots illustrated a similar time-dependent expression of *NaPAL4* transcripts in clusters 0, 3, and 4 (Fig. 7B). The transcript levels of the other BA biosynthetic genes in each cluster also depended on time with slight differences. *Na4CL1* and *NaAER1* expression peaked at ZT 12 in all clusters, but *NaPKS2* expression increased from ZT 8 to ZT 16 in the epidermal clusters, not in the other clusters. Next, we measured transcript levels of BA biosynthetic genes in corolla limb and tube (Fig. 7C). Although transcript levels of *NaPAL4* in the corolla limb did not differ with that in the corolla tube, *Na4CL1* expression was lower in corolla tube than in corolla limb. *NaPKS2* and *NaAER1* transcripts were barely detected in the corolla tube, which is consistent with the known location of BA biosynthesis (Kessler and Baldwin, 2007; Euler and Baldwin, 1996).

**Figure 7.**
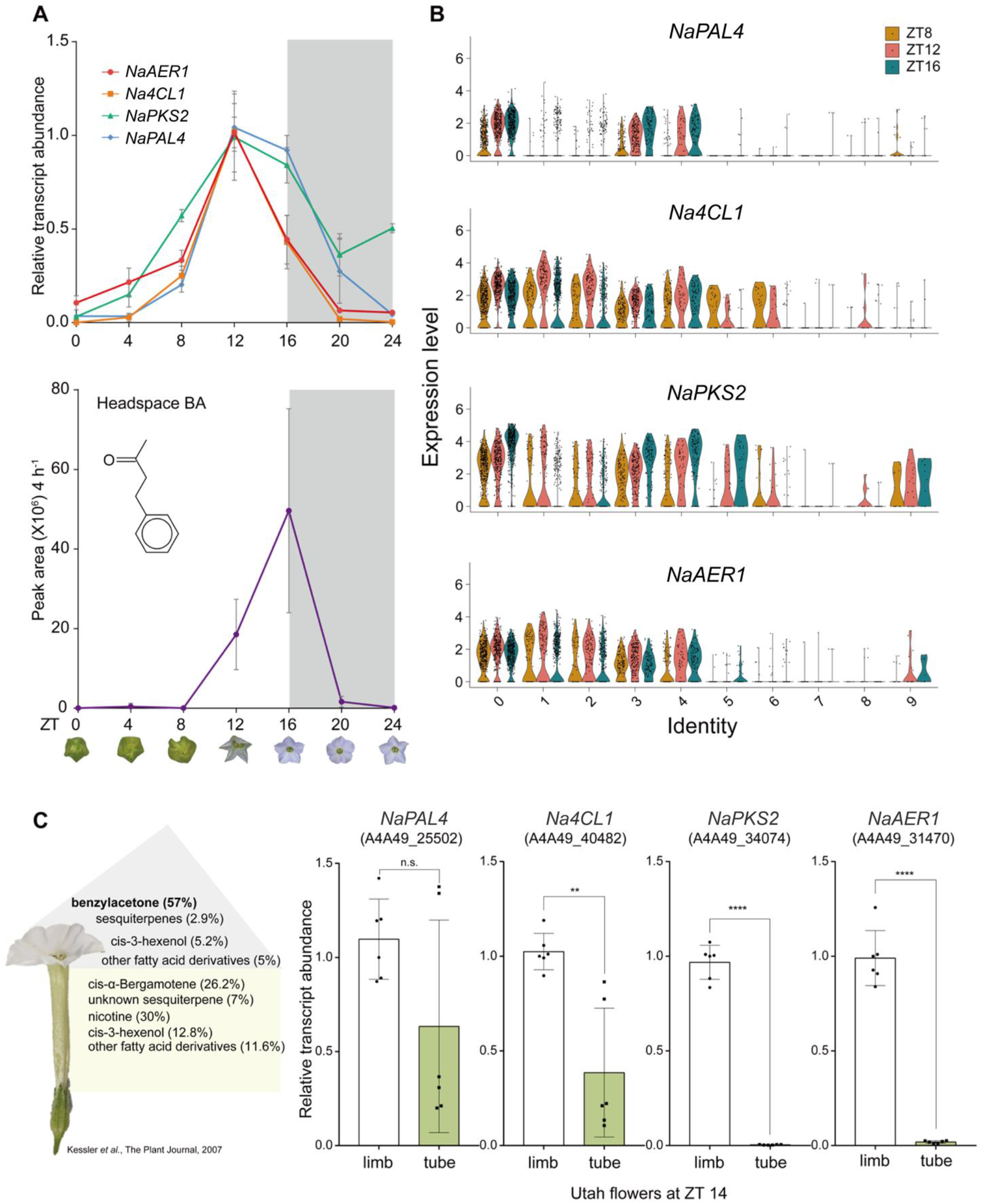
Diurnal rhythm of benzylacetone biosynthesis genes and its expression pattern in flower tissues. (A) Rhythmic expression of benzylacetone (BA) and BA biosynthetic genes. Relative transcript levels (mean ± SD) of *Na4CL1, NaAER1*, and *NaPKS2* peak at ZT 12, 4 hours before the peak of BA emission. (B) Violin plots show the expression levels of *NaPAL4, Na4CL1, NaPKS2*, and *NaAER1* across all clusters at three different time points. The level of *Na4CL1* and *NaAER1* transcripts peaks at ZT 12, as shown in Fig. 4a. *NaPAL4* and *NaPKS2* transcript levels at ZT 16 are higher than those at ZT 8 and 12 in cluster 0, 3, and 4. (C) Kessler *et al*. (2007) reported the percentage of *N. attenuata* floral scents in the floral headspace. BA is synthesized mainly in the corolla limb, and *cis*-α-bergamotene is synthesized in the corolla tube. Quantitative transcript analysis of BA biosynthetic genes in dissected flower tissues: limb and tube, reveals that the lack of BA emission in the corolla tube resulted from the limited expression of *NaPKS2* and *NaAER1* (mean ± SD; **, *p*<0.01; ****, *p*<0.0001; two-tailed Student’s t-test).

## Discussion

Transcriptional heterogeneity, both in different plant tissues and amongst accessions with distinctive floral volatile productions, has been used to identify key enzymes for volatile biosynthesis. Here, by analyzing the heterogeneity of cells in scRNA-seq of petal, we can prioritize candidate genes involved in floral scent biosynthesis. We also show that spatiotemporal expression of candidate genes provides reliable basis for selecting candidate genes. Bulk RNA-seq analysis has been used to examine gene-to-gene correlation, which helps to identify several key enzymes involved in floral scent biosynthesis (Xu et al., 2018; Boachon et al., 2019, 2015). We were also able to identify a CoA ligase in BA biosynthesis by calculating correlation coefficient among bulk RNA-seq transcriptome profiles. In this study, we can prioritize a PKS and a reductase involved in BA biosynthesis by performing the correlation analysis with the scRNA-seq dataset. Although *NaIFR3* expression was shown to be highly correlated with *NaPAL4* expression in the bulk RNA-seq dataset, we found that *NaAER1* was more correlated with *NaPAL4* than did *NaIFR3* in the scRNA-seq dataset. Why was *NaIFR3* expression strongly correlated with *NaPAL4* expression in the bulk RNA-seq? Members of an IFR family are known to bind a flavonoid which has a C6-C3-C6 carbon skeleton (Wang et al., 2006; Cheng et al., 2015). NaIFR3 may be able to reduce a flavonoid that originates from *t*-cinnamic acid (a NaPAL4 product) in other plant tissues.

BA is synthesized mainly in epidermal cells, as revealed from how cell populations clustered and from the spatial distribution of BA biosynthetic genes. NaPAL4, with its homologues NaPAL2 and NaPAL3, was found in the epidermis. Most cells (67 %) expressing *NaPAL4*, the gene required for the initial step in BA biosynthesis, harbored all other BA biosynthetic genes, indicating that the single-cell transcriptome map of *NaPAL4* largely concurred with the map of BA-producing cells in the corolla limb. Only 45% of *Na4CL1*-, 50% of *NaPKS2*-, and 48% of *NaAER1*-expressing cells contained the entire set of BA biosynthetic genes. High levels of *NaPAL4* and *Na4CL1* transcripts were found in the corolla limb and in the corolla tube (Fig. 7C), although no BA is produced in that tube. In contrast, *NaPKS2* and *NaAER1* transcripts were detected in the corolla limb and not in the tube, suggesting that the expression of *NaPKS2* and *NaAER1* determine BA production at the sub-organ level. Phe biosynthetic genes were mainly found in the epidermal clusters, which suggests that, in petals, primary metabolism is specialized to support the high demands of secondary metabolite production.

The single-cell transcriptome atlas of plant tissue helps us to identify biosynthetic genes of tissue-specific metabolites, because scRNA-seq analysis produces a promising correlation coefficient among transcriptome profiles. Rather than retrieving the average transcript levels of genes in tissue, we can examine gene-to-gene correlation using the transcript levels of genes expressed in a single cell. In addition, scRNA-seq generates high-dimensional data, which may provide more reliable correlation among genes (Shinozaki et al., 2018). Bulk RNA-seq data contains tens of thousands of variables (gene number), but the number of samples is much smaller than the number of genes. However, the scRNA-seq data has thousands of variables (gene number) and the number of cells is also more than thousands. Although gene co-expression analysis of scRNA-seq data is limited by low sequencing depth per cell (dropout events) and a high level of technical noise, we show that scRNA-seq analysis can be used to investigate biosynthetic genes of a target metabolite. Assuming that the spatiotemporal expression patterns of each set of genes in the biosynthetic pathway of a target metabolite are similar at the single-cell level, candidate genes for BA biosynthesis were prioritized. In addition, although protein sequence of NaPKS2 shared a highly similarity with NaPKS3 and transcript levels of both genes were similar in petals, scRNA-seq analysis predicted that NaPKS2 is the key enzyme for BA biosynthesis. scRNA-seq offers an alternative opportunity to investigate metabolic pathways in a single cell; such single-cell metabolisms are required for understanding secondary metabolite diversity within a plant.

## Methods

### Plant materials and protoplasting

Seeds of Utah 39^th^ inbred line were germinated on Petri dishes with Gamborg’s B5 medium including vitamins (Duchefa Biochemie, NL), as described previously (Krügel et al., 2002). Plants were grown under 16 h light/8 h dark conditions at 26°C with ± 2°C variation. Protoplasts were isolated from approximately 60 corolla limbs of *N. attenuata* at ZT 8, ZT 12, and ZT 16 of the first day of flower opening. Corolla limbs were excised by scalpel and immediately immersed in a plant culture dish containing 25 mL of cell-wall-degrading enzyme mixtures (Yoo et al., 2007). After 5 minutes of vacuum treatment, tissues were shaken at 20 rpm on an orbital shaker for 30 min in the dark, and the cell suspension was filtered onto a Petri dish using a 40 μm cell strainer (Corning, USA). The protoplast suspensions were centrifuged at 100 g for 5 min, and the resulting pellets were resuspended and washed with 10 % (w/v) mannitol solution. Cells were washed, centrifuged again, and finally filtered with 40 μm Flowmi tip strainer (Bel-Art SP Scienceware, USA). Cell concentration and viability were calculated using 0.2 % trypan blue on Cellometer Auto 2000 (Nexcelom Bioscience, USA). All samples showed a viability over 70 %. The entire protoplasting procedure was completed in 2 h.

### Generation of single-cell RNA sequencing library and feature-barcode matrices

Cell suspensions were loaded into the 10X Genomics Chromium single-cell microfluidics device with the Single Cell 3’ Library & Gel Bead Kit v3 (10× Genomics, USA) according to the manufacturer’s instructions. 12 PCR cycles were used for cDNA amplification, and 14 to 15 PCR cycles were performed for ligating adaptors. The library quality and size were checked using DNA high-sensitivity D5000 ScreenTape System (Agilent Technologies, USA). The single-cell RNA sequencing (scRNA-seq) libraries were sequenced on the HiSeq X Ten platform (Illumina, USA) at Macrogen (Republic of Korea). Approximately 400,000 reads and 1,300 genes were detected per cell.

Sequencing data were demultiplexed with mkfastq command in Cell Ranger (v.3.1.0) (10X Genomics) to generate fastq files. Expression matrices were also generated through the Cell Ranger pipeline, by aligning reads to the *N. attenuata* genome (NIATTr2). The Cellranger mkref command was used after filtering the GTF file to include only protein-coding genes in the *N. attenuata* genome. Feature-barcode matrix was generated after preprocessing, aligning, counting UMI, and filtering cells by Cell Ranger count command with default options.

Single-cell expression matrices were row-summed to generate pseudo-bulk datasets from scRNA-seq datasets. Conventional bulk RNA sequencing was performed with corolla limb at three different time points: ZT 8, ZT 12, and ZT 16. Pearson’s correlation coefficient was calculated between these pseudo-bulk datasets and conventional bulk RNA sequencing datasets. Some of the stress-related genes were upregulated in three scRNA seq datasets; these genes (described in Table S1) were excluded from downstream analysis.

### UMAP visualization and marker gene identification

After filtering matrices, the Seurat R package (v3.2.0) was used for dimension reduction and visualization (Satija et al., 2015; Butler et al., 2018). Cells that expressed < 200 genes were discarded. To integrate the single-cell libraries generated from three different time points, the datasets were normalized separately by SCT, and all features were selected for downstream integration. Pearson residuals were calculated by running PrepSCTIntegration, and anchors were identified by SCT normalization. Principal components analysis was conducted on the integrated dataset. Based on the top 50 principal components, distinct cell populations were defined by unsupervised clustering analysis (0.4 resolution) and visualized by uniform manifold approximation and projection (UMAP).

Marker genes of each cluster through all time points were selected by Wilcoxon rank sum test in Seurat’s FindConservedMarkers function. Genes over 0.25 log-fold change relative to other clusters and Bonferroni-corrected *p*-value <0.05 were chosen as upregulated genes in individual clusters. Average levels of transcripts and percentages of cells expressing the marker per cluster were visualized by color and size in dot plots or in violin plots. Based on the predicted functions of marker genes, we characterized clusters according to cell type, as epidermis, parenchyma, chlorenchyma, pollen, and vascular.

### Gene co-expression analysis

20 bulk RNA-seq datasets containing 14 tissue types from *N. attenuata* were downloaded from the sequence read archive database (Li et al., 2016). TPM values of each gene were used for calculating Spearman’s correlation coefficients between genes, and coefficients of each gene with *N. attenuata PAL4* (A4A49_25502) were extracted and visualized as histograms. For scRNA seq co-expression analysis, UMI values of genes were divided by the sum of UMI in each cell and multiplied by 100,000 to normalize the values. The normalized UMIs of each gene were used for calculating Spearman’s correlation coefficients between genes.

Genes with at least one UMI were assigned a value of 1, and others with no UMIs were assigned a value of 0. Cells with the value 1 of target gene were regarded as the cells expressing the gene. This binary matrix was then used as an input for an UpSet plot. The upset plot was drawn in ComplexUpset package (v.0.7.3) (Michał Krassowski, 2020; Lex et al., 2014). For visualizing the cells that express four BA biosynthetic genes, the FeaturePlot function in the Seurat package was used with the parameters: min.cutoff =“q10”, max.cutoff = “q90”, and blend = T.

### Virus-induced gene silencing

To silence a target gene in *N. attenuata* flowers, virus-induced gene silencing (VIGS) using tobacco rattle virus (TRV)-based vectors was performed (Saedler and Baldwin, 2004). A 250 bp fragment of the gene of interest was designed using the SGN VIGS Tool (Fernandez-Pozo et al., 2015) and cloned into the TRV2 vector. The *Agrobacterium tumefaciens* AGL1 strain harboring the TRV2 target gene was mixed with *Agrobacterium* containing TRV1 just before infiltration. About 1 to 2 mL of *Agrobacterium* solution was injected into *N. attenuata* rosette leaves. The infiltrated plants were incubated in the dark at 22°C for 2 days and then acclimated under weak light conditions in a growth chamber for 2 days. Later, the infiltrated plants were grown under 16 h light/8 h dark conditions at 26°C with ± 2°C variation. TRV2-eGFP was used as a negative control, and TRV2-PDS was used to observe VIGS efficiency. The silencing efficiency of individual flowers was measured by quantitative RT-PCR (qRT-PCR). Total RNA was extracted using RNeasy Plant Mini Kit (Qiagen, DE), and genomic DNA was removed by treating RNase-free DNase I (Qiagen). Extracted RNA was reverse-transcribed using oligo dT and SuperScript™ IV reverse transcriptase (Thermo Fisher Scientific, USA) following manufacturer’s protocols. Quantitative RT-PCR was performed on AriaMx real-time PCR System (Agilent technologies, USA) with TB Green Premix EX Taq (Takara, JP). Transcript levels were quantified by the comparative C_T_ method (Schmittgen and Livak, 2008). The Cq values were normalized to the value of the internal control gene, *EF1a-like*. Primers used in this study are listed in Table S2.

### Headspace and internal volatile sampling and GC-MS analysis

Flower headspace volatiles were trapped in silicone tubing (ST) and analyzed as described in Kallenbach *et al*. (Kallenbach et al., 2014). Briefly, before flowers start to open, a flower was excised and enclosed in a clean 8 mL glass vial with 300 μL of ultrapure water. Clean ST, 1 cm in length, was placed directly over the corolla limb for 3 h (ZT 14 – ZT 17). GCMS-QP2020 (Shimadzu, Japan) connected with a TD-30 thermal desorption unit (Shimadzu) was used for gas analysis. All standard compounds used in this study were purchased from Sigma-Aldrich (USA).

For internal volatile sampling, 50 mg of corolla limbs was finely ground in liquid nitrogen and resuspended in 1 mL of saturated calcium chloride solution spiked with tetralin as an internal standard (Joo et al., 2019). A 1 cm length of clean ST was incubated in the resuspension solution overnight, shaken at 80 rpm. Incubated STs were rinsed with ultrapure water and dried under a gentle nitrogen flow. Room temperature-equilibrated STs were analyzed in GCMS-QP2020 connected with a TD-30 thermal desorption unit.

### Functional evaluation of BA biosynthetic enzymes

Codon-optimized BA biosynthetic gene sequences (Bioneer, Republic of Korea) or native CDS sequences were cloned into a pET50 vector. *Escherichia coli* Rosetta (DE3) cells were transformed with pET50-a target gene and grown in LB broth media containing 50 µg/mL of kanamycin and 30 µg/mL of chloramphenicol at 37°C. When the OD_600_ reached 0.5 - 0.6, protein expression was induced by adding 0.35 mM of IPTG. After incubating an additional 18 - 20 h at 18°C, cells were harvested and lysed in cell resuspension buffer supplemented with 0.5 mM PMSF by sonication. Cell resuspension buffers for each proteins were prepared as previously described (Klempien et al., 2012; Koeduka et al., 2011; Abe et al., 2001): for purifying recombinant proteins, soluble protein fractions were obtained by centrifugation at 13,000 rpm for 20 min at 4°C, and loaded onto Ni-NTA agarose resin (Thermo Fisher Scientific), following manufacturer’s instructions. The size and purity of eluted proteins were confirmed by Coomassie staining. To get rid of excess imidazole, proteins were dialyzed against the final dissolving buffers and then stored at -80°C.

The CoA ligase activity of Na4CL1 with *t*-cinnamic acid was determined spectrophotometrically, as previously described (Klempien et al., 2012). The spectrophotometric assay was performed by monitoring the increase in absorption maximum at 311 nm for cinnamoyl-CoA production on a microplate reader. The assay reaction was performed with 250 µL solution containing 1.76 µg of purified Na4CL1 protein in 50 mM Tris/K phosphate/Na citrate pH 8.5, 0.05 mM *t*-cinnamic acid, 2.5 mM ATP, 5 mM MgCl_2_, and 0.4 mM CoA. Reactions were started by adding *t*-cinnamic acid to each well. 22,000 mL mmol^-1^cm^-1^ was the extinction coefficient used for calculating cinnamoyl-CoA formation (Lee et al., 1997).

We purified NaPKS2 and NaPKS3 proteins as described previously (Abe et al., 2001) and mixed NaPKS2 or NaPKS3 with the products formed during Na4CL1 spectrophotometric assays and malonyl-CoA. The reaction mixture contained 50 µL of Na4CL1 products (intended to contain final 40 µM of cinnamoyl-CoA in 500 µL reaction mixture), 40 µM malonyl-CoA, and 10 µg of NaPKS2 or NaPKS3 in a final volume of 500 µL of 100 mM potassium phosphate buffer containing 1 mM EDTA (pH 8.0). Mixtures were incubated at 30°C for 2 h, and reactions were stopped by adding 50 µL of glacial acetic acid. The products were extracted with 1 mL of ethyl acetate via vigorous vortexing, and the ethyl acetate layer was concentrated using a freeze-dryer. All products were injected into GC-MS via thermal desorption unit (Shimadzu).

Double-bond reducing activity of NaAER1 was determined as described previously (Koeduka et al., 2011). The assay mixture contained 3 µg of purified NaAER1 proteins, 0.5 mM NADPH, 0.5 mM benzalacetone in a final volume of 400 µL of 125 mM of MES-KOH (pH 6.5). Incubations were carried out for 20 min at room temperature, and the reactions were stopped by adding 15 µL of glacial acetic acid. The products were extracted with 1 mL of ethyl acetate and concentrated. Products were injected into GC-MS via TD unit.

## Acknowledgments

We appreciate Dr. Inkyung Jung, Seongwan Park, and Mooyoung Kim for assisting us to generate scRNA seq library; Dr. Gisuk Lee and Dr. Yongsung Joo for help with metabolite analysis; San-Hae Lim with protein purification; Guo Han, Shuqing Xu, and Ian T. Baldwin for comments on BA biosynthetic pathway; Emily Wheeler for editorial assistance. This work is supported by grants from the Samsung Science & Technology Foundation (SSTF-BA1901-10, S.-G.K.) and TJ Park Doctoral Science Fellowship (M.K.).

## Author contributions

S.-G.K. conceived and supervised the project. M.K. generated scRNA-seq libraries and performed bioinformatic analysis. Y.C. generated cDNA libraries for bulk RNA sequencing. M.K. and H.K. analyzed plant volatiles. M.K. and H.K. performed VIGS experiments. M.K. conducted quantitative PCR analysis and *in vitro* enzyme assay. S.-G.K. and M.K. wrote the manuscript.

## Additional information

Supplementary information will be available for this paper.

## Data availability

scRNA-seq and bulk RNA-seq libraries will be deposited after acceptance. All other data generated during this study are included in this published article and supplementary information files.

